# Alternative Growth Behavior of *Mycobacterium Avium* Subspecies and Staphylococci with Implications for Clinical Microbiology and Blood Culture

**DOI:** 10.1101/049031

**Authors:** Peilin Zhang, Lawrence M. Minardi, J. Todd Kuenstner, Steve M. Zekan

## Abstract

Rapid culture of *Mycobacterium* avium subspecies paratuberculosis (MAP) from patients remains a challenge. During the process of developing a rapid culture method for MAP, we found that there is an alternative growth behavior present in MAP, MAH (*Mycobacterium avium* subspecies hominissuis) and other bacteria such as *Staphylococcus aureus*, and *Staphylococcus pseudintermedius*. The bacterial DNA, RNA and proteins are present in the supernatants of the liquid culture media after routine microcentrifugation. When cultured in the solid media plate, there are a limited number of colonies developed for MAP and MAH disproportionate to the growth. We believe there is an alternative growth behavior for MAP, MAH and other bacteria similar to “phenoptosis”. Based on the alternative bacterial growth behavior, we tested 62 blood culture specimens that have been reported negative by routine automated blood culture method after 5 days of incubation. We used alternative culture media and molecular diagnostic techniques to test these negative culture bottles, and we found a large percentage of bacterial growth by alternative culture media (32%) and by molecular PCR amplification using 16s rDNA primer set and DNA sequencing (69%). The sensitivity of detection by the molecular PCR/sequencing method is significantly higher than by routine automated blood culture. Given the challenge of early diagnosis of sepsis in the hospital setting, it is necessary to develop more sensitive and faster diagnostic tools to guide clinical practice and improve the outcome of sepsis management.

## Introduction

*Mycobacterium avium* subspecies *paratuberculosis* (MAP) is known to be a slow grower with a special requirement for mycobactin as a growth factor in the culture media (1, 2). MAP has been extensively studied in Johne’s disease, a veterinary condition in cattle and sheep, and it has been associated with Crohn’s disease (CD), although the association is controversial (2). The technical difficulty of growing MAP in culture lead us to search for an alternative method to grow MAP more rapidly to determine the relationship of MAP and CD and to multiple other potential conditions such as multiple sclerosis, and type I diabetes mellitus (3-5). In the natural environment where MAP is present, dependence of MAP on mycobactin for growth is likely to be provided by another Mycobacterium to produce mycobactin if the dependence is absolute, and experimentally it has been shown that the mycobactin dependence of MAP is not absolute (1, 6, 7). Even in the presence of mycobactin, MAP growth in solid media or liquid media requires an extended period of time (up to 1 years), and this prolonged culture process makes it mathematically unimaginable, since it becomes more than a matter of bacterial cell division (one cell divides to two, two cells divide to four, and so forth) and duplication. This slow growth pattern of MAP leads us to believe there is a destructive force during MAP culture. The best known mechanism of a destructive growth pattern is the transiting of bacteriophage from pro-phage to lytic cycle of mature phage during growth of *E. coli* (8). The phenomena of the lytic cycle of *E. coli* growth with bacteriophage have been described as a specific type of programmed cell death (phenoptosis)(9). MAP phages (Mycobacterium phage vB_MapS_FF47, Genbank # NC_021063) have been previously identified and isolated without known implication or functions (10). Another mycobacteriophage with highly homologous DNA sequence to phage FF47, phage Muddy, had also been isolated from the soil samples from Africa (Genbank # NC_022054). There are other mycobacteriophages known to infect MAP as well as other Mycobacterial species (11). Significant effort was made to determine the presence of phage DNA by PCR and next generation sequencing from our lab to no avail.

*Mycobacterium avium* subspecies *hominissuis* (MAH) is one of the closest members within the subfamily of *Mycobacterium avium* subspecies with significant homology in structure and functionalities to MAP. However, MAH is a much faster grower with significant clinical implications in immune compromised patients such as cancer patients undergoing chemotherapy and HIV/AIDS patients (12, 13). We have previously isolated a strain of MAH, MAH-ashley, from a Crohn’s patient (14). MAH-ashley grows fast with colonies in 24-48 hours on the solid media plate. We also demonstrated a similar growth pattern of MAH-ashley to that of MAP. We used *Staphylococcus aureus* and *Staphylococcus pseudintermedius* as controls to show similar growth in liquid culture are present in other bacteria. Due to the specific bacterial growth in liquid media, automated blood culture instruments in the clinical laboratory may or may not be able to detect the presence of bacterial growth. This at least partially explains why the yield of routine blood culture is so low and the diagnosis of sepsis in hospital patients is often empirical at best.

## Materials and methods

### 1, Bacterial culture in liquid and solid media

MAP Dominic strain (ATCC Cat: 43545) was purchased from ATCC and maintained in the modified liquid and solid media plate described previously (14). MAH was isolated from the blood of a Crohn’s patient by our laboratory previously and was maintained in liquid and solid media as previously described (14). MAH isolated from our lab, MAH-ashley, has been further characterized by full genomic sequencing (Next generation sequencing, Illumina Miseq Sequencing System) from West Virginia University (WVU) Genomics Core facility, Morgantown, WV, and the sequencing data were analyzed and compared with the existing MAH and other Mycobacterial sequences in Genbank (unpublished data). MAH-ashley is a fast grower with visible colonies on the solid media plate within 24-48 hours. Three different types of media, liquid media, regular solid media plate, and induction media were routinely used for culture of MAP and MAH (14). *Staphylococcus aureus* was isolated from the blood of a chronically ill patient through induction culture method based on the Middlebrook 7H10 media, and this isolate can grow in all culture media including Brain-heart infusion broth/plate, Tryptic Soy broth/plates as well as chocolate agar plate. Partial genomic DNA sequencing using 16s rDNA primers and subsequent full genomic sequencing (NGS) by WVU Genomics Core Facility were also performed for characterization of the isolate (unpublished data). *Staphylococcus pseudintermedius* was isolated from the skin wound of a domestic dog, and the growth behavior of this isolate is similar to that of *S. aureus*. Similar partial genomic sequencing and full genomic sequencing by NGS were also performed on this isolate of *S. Pseudintermedius* for confirmation (unpublished data). All culture media components were purchased from BD Biosciences (Middlebrooke 7H9, 7H10, Tryptic Soy media, Brain-heart infusion media, media additives), and prepared in our lab according to the manufacturer’s instruction. Mycobactin J was obtained from Allied Monitor Inc., MO.

### 2, DNA, RNA and protein extraction and gel electrophoresis

All DNA, RNA and protein isolation was carried out as described previously using standard methods except for pre-treatment of the bacterial/mycobacterial cultures with acetone (8, 14). DNA and RNA were visualized on the agarose gel electrophoresis for integrity and visual estimation. All proteins were visualized on SDS-polyacrylamide gel electrophoresis.

### 3, PCR analysis of bacterial culture and Mycobacterial culture

PCR amplification of specific genes for MAP, MAH and 16s rDNA was performed as described previously (14). Partial genomic sequencing of PCR amplicons was performed at Eurofins Genomics facility in Louisville, KY. The partial DNA sequences generated from the PCR amplicon sequencing were compared against Genbank using the existing BLAST nucleotide search engine at NCBI (http://blast.ncbi.nlm.nih.gov/Blast.cgi).

### 4, Routine automated blood culture and subsequent analyses

Routine automated blood culture bottles from bioMerieux BacT/Alert 3D clinical system were obtained from the local hospital after 5 days incubation, and these blood culture bottles were reported as “negative” and to be discarded. We only tested aerobic culture bottles as a proof of principle, and no anaerobic or fungal cultures were examined. After a proper IRB expedited review approval, the culture bottles were open in the level 2 biosafety cabinet hood, and two 200 ml aliquots were removed: one aliquot was used for alternative induction culture and another aliquot was for DNA/PCR analysis. Cultures on the alternative media slope were kept for 72 hours, and the results were collected/recorded. A longer incubation time was not tested. The alternative induction culture medium was developed in our laboratory as described previously with the base media Middlebrook 7H10 agar supplemented with yeast extract, glycerol and 20% sucrose (14). The culture media were prepared in our laboratory and autoclaved for 121°C for 20 minutes with pressure of 21 psi (standard liquid cycle). An aliquot of 200 ml was taken from the culture bottle and directly inoculated on the slope of alternative culture media. The inoculated blood culture on the slopes were incubated in a 37°C incubator for 72 hours, and examined by direct visualization of colonies on the slopes. Visible colonies on the slopes are reported as positive.

An additional 200 ml blood culture was taken from the blood culture bottle, and transferred to a sterile Eppendorf micro-centrifuge tube. The blood culture is centrifuged at 12000g for 5 minutes to separate the cell pellet/debris from the supernatant. An aliquot of 20 ml of the supernatant was directly loaded on 0.8% agarose gel for electrophoresis. The cell pellet is washed once with 100% acetone, and resuspended in 200 ml TE buffer (pH 7.6) before being heated at 95°C for 15 minutes. The aliquot is centrifuged at 12000 g for 2 minutes, and 0.5 ml of the supernatant is directly used for PCR amplification in 25 ml total volume. PCR primers were based on the 16s rDNA primer set described previously (14). PCR amplification was visualized on 6% non-denaturing polyacrylamide gel or 1.2% agarose gel electrophoresis, and the PCR amplicons were stained with ethidium bromide (8). Direct digital photographs were taken. PCR amplicons were sent to Eurofins Genomics services, Louisville, KY for direct sequencing and the DNA sequences were subjected to BLAST search of Genbank at NCBI.

## Results

### 1, Alternative growth behavior of MAP, MAH, *S. aureus* and *S. pseudintermedius* in the liquid culture media (supernatant)

The fact that the growth of MAP in vitro takes several months leads us to believe there is an alternative pathway, and the same could be true for *Mycobacterium* in general *including Mycobacterium tuberculosis* which requires up to six weeks for growth. The best characterized molecular mechanism for bacterial lytic culture is through bacteriophage. We sought to identify if there are mycobacteriophages within MAP and MAH-ashley. We maintained the culture in liquid media for MAP, MAH-ashley, *S. aureus* and *S. pseudiointermedius* in their respective culture media suitable for growth. MAP and MAH-ashley were maintained for 3-weeks and 3 days respectively in Middlebrook 7H9 broth with various supplements, and *S. aureus* and *S. pseudintermedius* were cultured in the brain-heart infusion (BHI) broth for 48 hours. One milliliter of liquid culture from each tube was removed and centrifuged for 10 minutes at 15000 g in microfuge. The supernatants were transferred to fresh microfuge tubes, and 20 ml was loaded on 0.7% agarose gel for electrophoresis with ethidium bromide (Figure 1A). There are clear bands over 10 kb in size in all the culture tubes, indicating the presence of specific DNA fragments in the culture media supernatants. The fact that there are DNA fragments in specific size suggests the presence of a bacteriophage. However, examination of the DNA isolated from the supernatants of MAP and MAH cultures by NGS twice failed to demonstrate the presence of any phage DNA (unpublished). Similarly 20 ml aliquots of culture media supernatants were loaded on 1.2% agarose formaldehyde gel using MOP buffer (Figure 1B), and there are specific RNA within all the culture media. Aliquots of 20 ml culture media supernatants were loaded on 4%/10% SDS-polyacrylamide gel for protein analysis and stained with Coomassie blue (Figure 1C). There are clearly multiple bacterial proteins within the supernatants of all cultures after the centrifugation, in addition to DNA and RNA (Figure 1A and 1B). There were more proteins in the supernatants of MAP and MAH cultures than the cell pellet lysate (Figure 1 C). These results indicate MAP and MAH growth and lysis occur simultaneously with various speeds. MAP, being the slow grower, cannot generate more MAP cell pellet in the extended culture period, whereas MAH, the fast grower, generating much larger cell pellet, also produces extracellular contents in the supernatant through an unknown process. Given the specific DNA, RNA and protein contents within the culture media, we felt this process is likely a specific programmed cell death similar to “phenoptosis” described previously (9).

**Figure 1:**
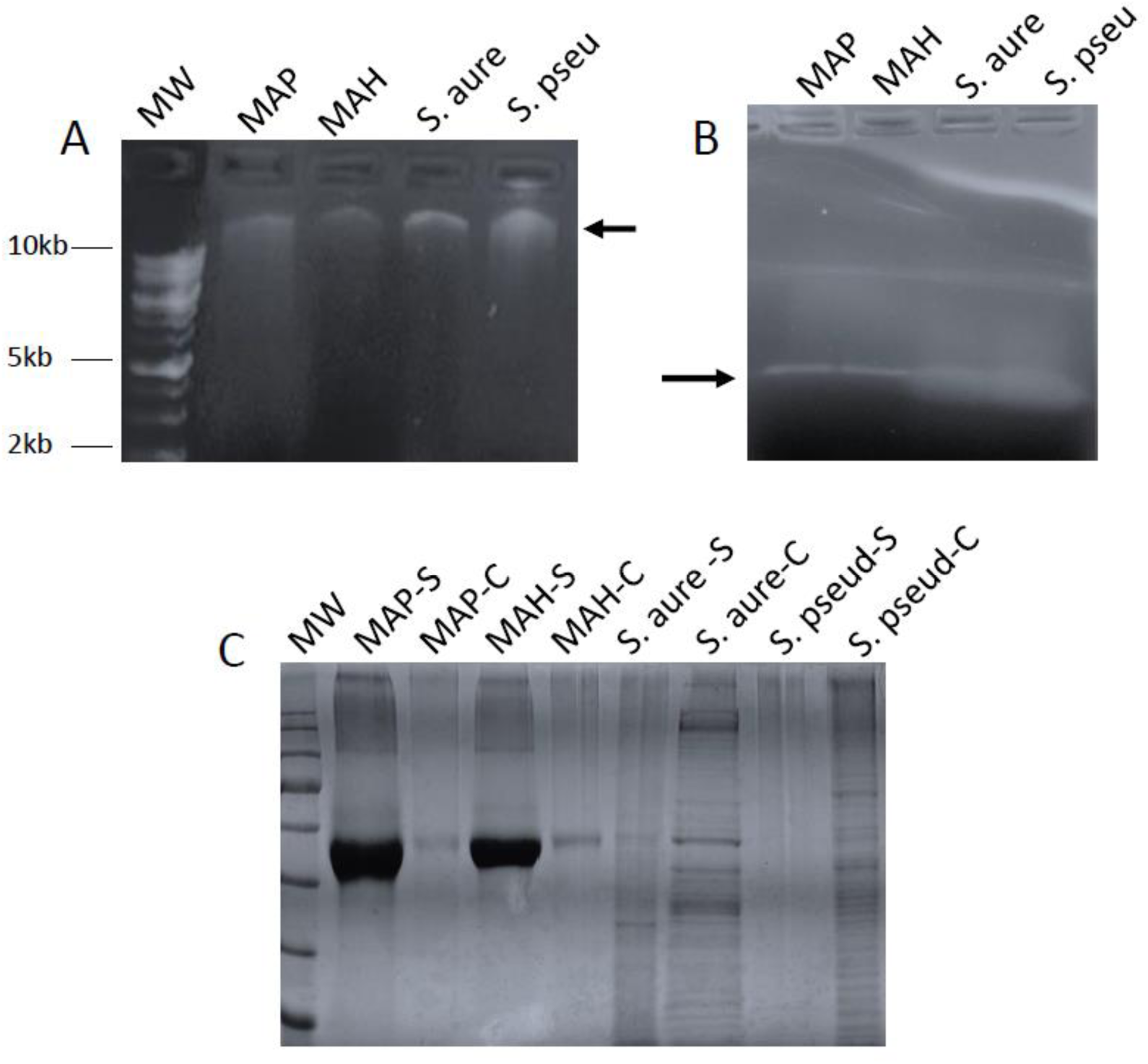
The presence of DNA, RNA and proteins in the culture media of MAP, MAH, S. aureus, and S. pseudintermedius. A: 0.7% agarose gel with ethidium bromide. MW is 1kb molecular weight marker with maximal 10 kb. B: 1.2% agarose formaldehyde gel with ethidium bromide. C: 10% SDS-PAGE. Each gel lane contains 20 μl culture media directly loaded on the gel after centrifugation. The high intensity band in MAP-S and MAH-S represents added albumin of the OADC supplement in the 7H9 culture media.

### 2. Bacterial growth pattern and clinical automated blood culture

Based on the results of the alternative growth through the process of likely “phenoptosis” described above, we examined 62 blood culture bottles obtained from the local hospital using the bioMerieux BacT/Alert 3D automated blood culture system. We only examined the aerobic culture bottles for proof of principle. These blood culture bottles were to be discarded after 5-day incubation and the “negative” culture results had been reported clinically. Two separate aliquots of 200 ml each were obtained: one aliquot was inoculated on the slope of induction media as described previously, and one aliquot was used for DNA and PCR/sequencing analysis. The slope culture was kept for 72 hours at 37°C and the results of visible colonies were recorded. We only report the presence (positive) or absence (negative) of visible colonies on the media slopes, and we did not count the number of colonies to quantify the bacterial growth. The patients’ clinical and demographic information was not obtained since the study was designed to compare the culture and detection methods for bacterial growth. One aliquot of the culture was centrifuged for 5 minutes at 15,000 g in microfuge and the supernatants were directly used for 0.7% agarose gel electrophoresis (Figure 2 A). The cell pellets were treated with 100% acetone once, and then resuspended in 200 ml TE buffer (pH 7.6). The resuspended cell pellets were heated at 95°C for 15 minutes, and directly used for PCR analysis using the 16s rDNA primer set as described (Figure 2B)(14). The PCR amplicons were sent for sequencing analysis to Eurofins Genomic services. The BLAST results of the PCR amplicons are listed in Table I for the best match identification. As in Table I, there were 20 positive culture results with visible colonies on the induction media containing 20% sucrose (20/62, 32.2%), and 43 of the 62 culture bottles were positive by PCR using the 16s rDNA primer set as described (43/62, 69%). The bacteria identified by DNA sequencing of the PCR amplicons were diverse and listed as in the last column in Table 1.

**Figure 2:**
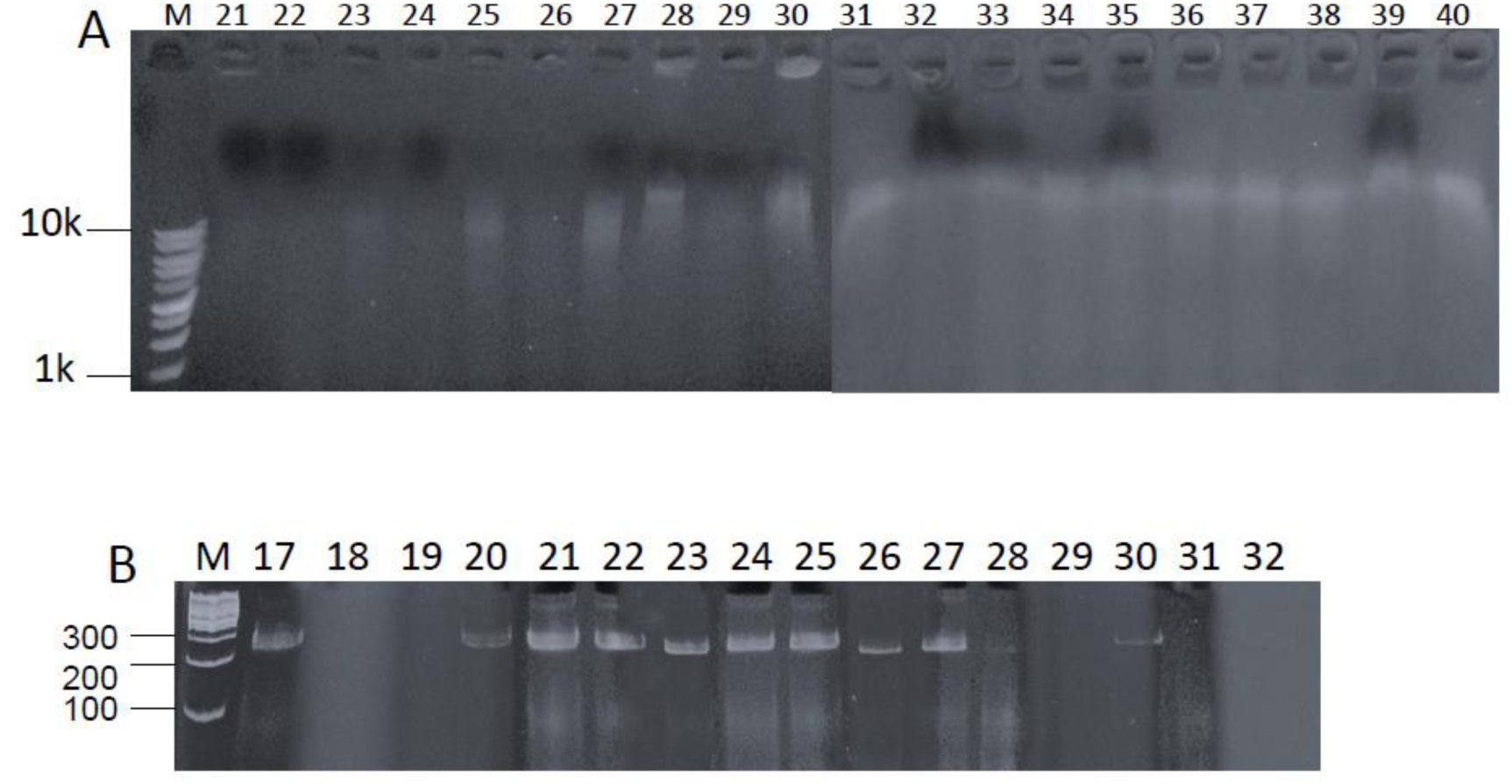
The presence of bacterial DNA within the culture media and PCR amplification from the automated blood culture system. A: 0.7% agarose gel electrophoresis of blood culture media after centrifugation. MW is the 1kb molecular weight marker. B: 6% non-denaturing acrylamide gel electrophoresis of PCR amplification using the DNA from the automated blood culture bottles. MW is the 100 bps molecular weight ladder.

**Table I:**
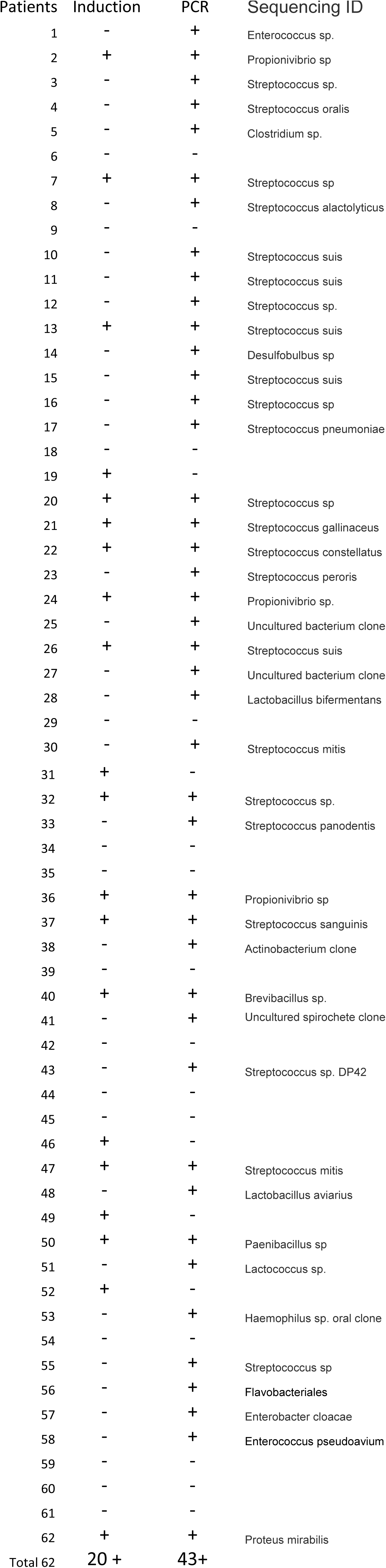
Culture on induction media slants and PCR/sequencing identification

## Discussion

MAP growth is an area of great interest in the diagnosis of CD in humans and Johne’s disease in cattle and sheep. The technical difficulties are multiple, and the established method for MAP culture is clinically impractical due to the prolonged period of incubation. This technical challenge makes it difficult to convincingly demonstrate the role in pathogenesis of Crohn’s disease. Assuming that the doubling time of MAP replication is 14-20 hours (1), a single cell may be estimated to yield a visible colony in less than 30 days. To incubate the culture for 6 months is difficult to imagine from the standpoint of basic cell division. This suggests that there is an alternative unknown growth behavior of MAP. The DNA, RNA and proteins present in the liquid media (extracellular) suggest a lytic growth pattern without a phage. We also noted the same growth patterns in MAH, *S. aureus* and *S. pseudintermedius*. We have identified a strain of *E. coli* from a patient with nephrotic syndrome with similar growth behavior. We also filtered the culture supernatant with 0.2 mM sterile syringe filter, and we didn’t see colonies on the plate after filtration (data not shown). The presence of bacterial extracellular DNA, RNA and proteins fits the description of “phenoptosis”. Protein-leaky mutants of *E.coli* have been previously described with chemical mutagenesis, and these leaky mutants were reported to form blebs on the cell surface without lysis of the bacteria(15, 16). But these protein-leaky mutants of *E.coli* reduced approximately 15% of the protein contents. No bacterial DNA leakage was reported or studied.

Given the growth behavior of MAP, MAH, *S. aureus* and *S. pseudintermedius* with likely phenoptosis in our study, we sought to see if similar growth behavior is present in the patients with routine automated blood culture. Clinically it is known that the yield of the automated blood culture system ranges from 5 to 15% in hospital patients. This low yield could be due in part to the bacterial growth such as phenoptosis. We used an alternative media with high osmolarity (induction media with 20% sucrose) for induction of specific growth, and this high osmolarity media increased the yield of culture significantly. The detailed mechanism of increased or decreased bacterial growth is unclear, and it appears the effect is species dependent. The presence of bacterial DNA by PCR/sequencing in clinically culture negative patients raises the question of whether automated routine blood culture systems have sufficient sensitivity. The indication for blood culture is clinically determined, and the diagnosis of sepsis in the hospital setting is a critical step for management. It is important to have an alternative detection method to compare with routine blood culture system. Given the presence of bacterial phenoptosis, rapid molecular diagnostic tools seem critical for the patients in hospital.

Recent description of bacterial DNA translocation is an interesting concept in CD and inflammatory bowel disease (IBD) (17, 18). It is also interesting that common pathogenic bacteria such as enterobacteria, Staphylococci, and Streptococci, etc., cannot be clinically cultured from the blood of IBD patients (17, 18). “Bacterial translocation” has been previously described that the epithelial barrier of intestinal mucosa is defective and the intestinal bacteria will travel through the barrier to the blood circulation and interact with inflammatory cells within the lamina propria or submucosa. The best example of bacterial translocation through epithelial cells and the lymphatic system is *Salmonella typhi* through cystic fibrosis transmembrane conductance regulator (CFTR) by the epithelial cell internalization (19-21). However, bacterial “naked DNA” translocation is unknown, and the bacterial proteins in the blood cannot be demonstrated unless specific antibodies against bacterial proteins are employed. The fact that up to 20-50% Crohn’s patients are positive for anti-ompC antibody indicates a host specific response to ompC of *E. coli* (22). The presence of bacterial DNA within the blood circulation is likely a representation of bacterial infection based on our study of MAP culture, and we felt the phenomena fit the description of “phenoptosis”. A type of phenoptosis was described for lytic cycle growth of *E.coli* with phage (9), and we were not able to demonstrate the presence of mycobacteriophage in MAP and MAH. More studies will be needed to elucidate the mechanism of this peculiar pattern of growth of MAP, MAH and other medically important bacteria.

